# Encoding of the Intent to Drink Alcohol by the Prefrontal Cortex is blunted in Rats with a Family History of Excessive Drinking

**DOI:** 10.1101/490664

**Authors:** David N. Linsenbardt, Nicholas M. Timme, Christopher C. Lapish

**Affiliations:** Addiction Neuroscience, Department of Psychology and Indiana Alcohol Research Center, Indiana University – Purdue University Indianapolis, Indianapolis, IN 46202

**Keywords:** alcohol-associated cues, alcohol-preferring rat, prefrontal cortex, electrophysiology, neural encoding, information theory, decision-making

## Abstract

The prefrontal cortex plays a central role in guiding decision-making, and its function is altered by alcohol use and an individual’s innate risk for excessive alcohol drinking. The primary goal of this work was to determine how neural activity in the prefrontal cortex guides the decision to drink. Towards this goal, the within-session changes in neural activity were measured from medial prefrontal cortex (mPFC) of rats performing a drinking procedure that allowed them to consume or abstain from alcohol in a self-paced manner. Recordings were obtained from rats that either lacked or expressed an innate risk for excessive alcohol intake - Wistar or Alcohol Preferring ‘P’ rats, respectively. Wistar rats exhibited patterns of neural activity consistent with the intention to drink or abstain from drinking, whereas these patterns were blunted or absent in P rats. Collectively, these data indicate that neural activity patterns in mPFC associated with the intention to drink alcohol are influenced by innate risk for excessive alcohol drinking. This observation may indicate a lack of control over the decision to drink by this otherwise well-validated supervisory brain region.

---

Aberrant decision-making is both a risk factor for, and the result of, an Alcohol Use Disorder (AUD; (Verdejo-Garcia et al., 2017). Therefore, understanding the neural systems that underlie decision-making, and how altered function of these systems influences decisions about drinking alcohol, is critical to identify novel targets to treat and prevent AUDs. While several neural systems have been implicated in decision-making, the medial prefrontal cortex (mPFC) plays a critical role in setting goals (Buschman and Miller, 2014) and forming intentions to achieve them (Fuster and Bressler, 2015, Brass et al., 2013, Haynes et al., 2007). Thus, the inability to refrain from excessive drinking may reflect pathology in neural circuits that guide goal-directed actions such as mPFC (Fuster and Bressler, 2015).

Dysfunction of the mPFC has been repeatedly found in populations of subjects that drink alcohol excessively (Schacht et al., 2013). Exposure to experience-or experimentally-paired alcohol cues, increases neuronal activity within the PFC (Tapert et al., 2003, George et al., 2001, Kareken et al., 2010), and the magnitude of this effect is correlated with increases in self-reported alcohol craving (Myrick et al., 2004) and relapse (Grusser et al., 2004). Additionally, recently abstinent individuals with an AUD exhibit reduced baseline neuronal activity within the mPFC (Catafau et al., 1999). Similar effects are observed in rodents, with exposure to alcohol-associated cues eliciting reinstatement of extinguished alcohol seeking and robust increases in biomarkers of neural activity in PFC (Dayas et al., 2007, Groblewski et al., 2012). More recent reports suggest a critical role for the PFC in alcohol extinction learning (Keistler et al., 2017, Cannady et al., 2017), suggesting that this brain region may be critically involved in ‘remapping’ associations between alcohol-associated stimuli and the motivational properties of alcohol. Thus, preclinical rodent and human data converge to implicate altered function of PFC in AUD.

The PFC has also long been known to be involved in the regulation of executive processes required to guide reward-based decision-making (Bechara, 2005, Ridderinkhof et al., 2004, Krawczyk, 2002), and animal studies are beginning to shed light on the computational processes that underlie these decisions (Dalley et al., 2004, Fitoussi et al., 2015). Decisions to initiate (or suppress) reward-directed motor actions are encoded in frontal-parietal circuits (Andersen and Cui, 2009), and, in the PFC, the encoding of these actions are evident prior to action initiation indicating behavioral intent (Sakagami and Niki, 1994, Sakagami and Tsutsui, 1999, Tanji and Hoshi, 2001, Momennejad and Haynes, 2013, Boulay et al., 2016, Andersen and Cui, 2009). These data motivated our hypothesis that similar neurocomputational processes exist in the PFC that regulate alcohol intake decisions. The implications of identifying and understanding processes that underlie the intention to use alcohol cannot be overstated, because intention signals that arise prior to alcohol seeking/drinking may be particularly effective targets for interventions aimed at reducing or eliminating alcohol consumption.

To first determine if the signals reflecting the intention to drink alcohol were present in the PFC, the current study evaluated neural activity across populations of neurons recorded during alcohol drinking in well-trained, high drinking, rats. We were particularly interested in the impact of alcohol-associated cues on drinking intent, and the role of family history of alcohol drinking on these cue-elicited decisions, as these factors have been shown to be critically important in human clinical studies (see above). Thus, we used Indiana alcohol-preferring ‘P’ rats, which are a well-validated preclinical model of transgenerational risk for excessive drinking (i.e. ‘family-history positive’), and a comparison strain with no family history, Wistar rats. We hypothesized that the intention to drink or abstain would be encoded in populations of neurons in the PFC. Furthermore, since individuals with a positive family history display greater PFC responses to alcohol associated stimuli (Kareken et al., 2010, Tapert et al., 2003), we also hypothesized that P rats would display a more robust intention signal compared to Wistar.

## Materials and Methods

### Animals

P rats have been selectively bred for > 75 generations for their high drinking phenotype (Bell et al., 2006, Li and McBride, 1995, McBride et al., 2014), and are conceptually analogous to individuals with generations of family history of excessive drinking (i.e. family history positive). As P rats were originally derived from Wistar rats, we opted to use this population (which is ‘family history negative’) to assess possible family history effects.

P rats were ordered from the Indiana Alcohol Research Center Animal Production Core (Indianapolis, IN), and Wistar rats were ordered from Envigo (Indianapolis, IN). All animals were shipped via truck to our vivarium, and were single housed and placed on a 12 hour reverse light/dark cycle. Animals were ≈70 days of age prior to testing and had *ad lib* access to food and water. All procedures were approved by the Purdue School of Science Animal Care and Use Committee and conformed to the Guidelines for the Care and Use of Mammals in Neuroscience and Behavioral Research (National Academic Press, 2003).

### Intermittent Alcohol Procedure (IAP)

The procedural timeline and methods for these experiments have been recently described in detail (Linsenbardt and Lapish, 2015). All animals first underwent an IAP using previously published procedures (Simms et al., 2008): Rats were given access to 2 bottles, one containing 20% alcohol (v/v) and the other tap water, for 24 hours every other day (Mon/Wed/Fri) in the home cage. These procedures were continued for 4 weeks; animals had 12 total 24-hour alcohol/water access sessions.

### 2-Way Cued Access Protocol (2CAP)

Twenty-four hours following the final (12^th^) IAP access session animals received access to an unsweetened 10% alcohol (v/v) solution for 2CAP sessions. 2CAP sessions occurred during the dark phase of the light/dark cycle, starting 1-3 hours after lights off. The conditioning box configuration is illustrated in Figure 1A. During 2CAP, a white stimulus light was illuminated for 2 seconds on one side of the rectangular box at random. One second after this light was turned off, a sipper tube containing 10% alcohol (v/v) solution was extended into the box on the same side as the light cue. Thus, the light was a Discriminative Stimulus (DS+) that predicted the location that the alcohol was to be made available. To ensure the sipper motor sound did not serve as a directional cue, both tube motors were turned on for the same duration, but only the appropriate sipper entered the chamber. The tube was available for ≈10 seconds. Each trial was separated by a 20-180 second inter-trial interval (ITI; 90 seconds on average; randomized order). A total of 40 trials were conducted for 5 out of 7 days a week (weekdays) for 3 weeks (15 total sessions) prior to surgery. Water sessions were identical to alcohol sessions except the sippers contained water. During water sessions, a tube containing 10% (v/v) alcohol was present outside the fluid delivery port to ensure that the presence or absence of the alcohol odor did not predict alcohol availability/unavailability.

**Figure 1.**
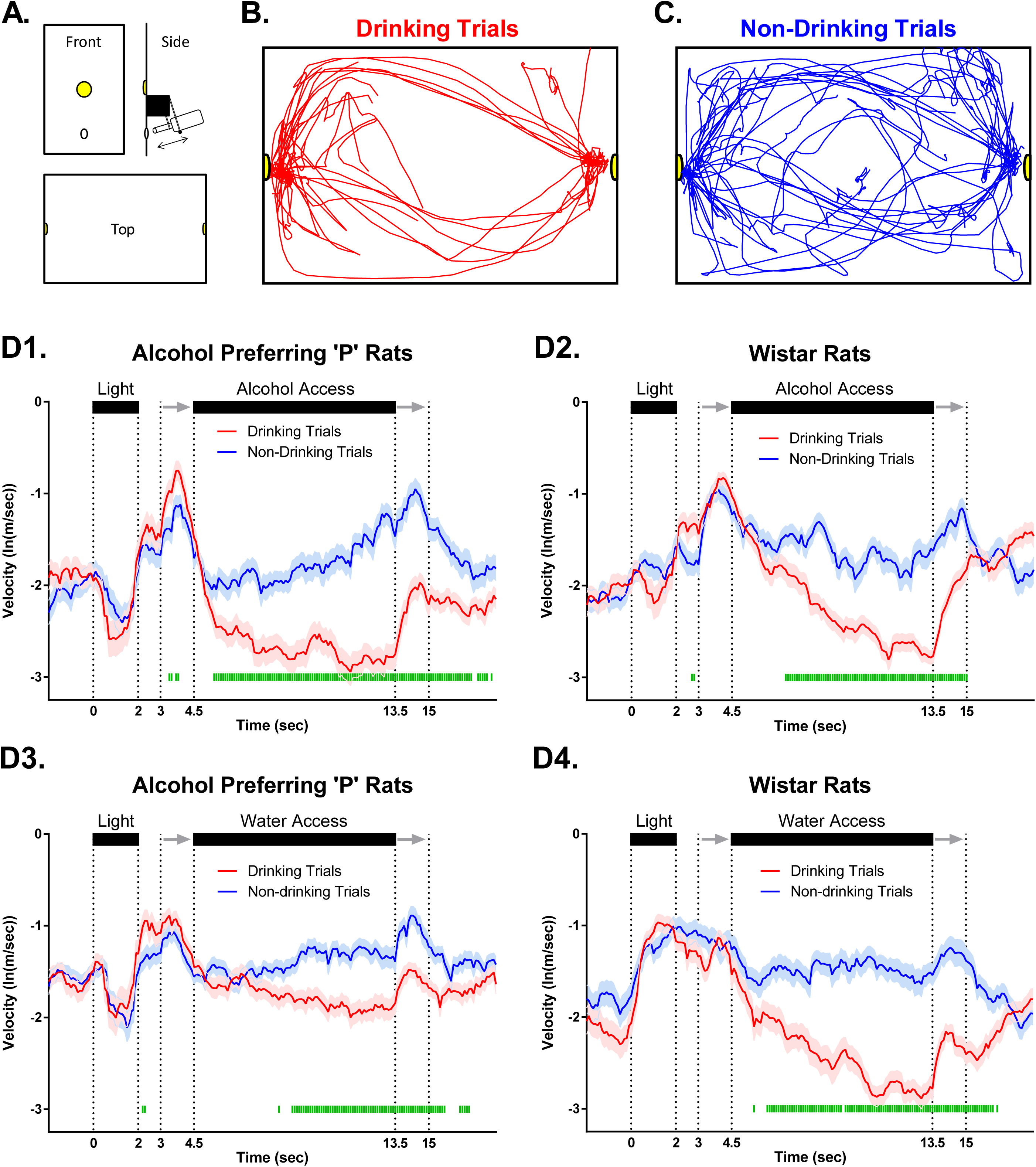
Movement dissociates drinking versus non-drinking trials during fluid availability but not during stimulus (Discriminative Stimulus; i.e. cue light) presentation. **(A)** Configuration of conditioning boxes used for cue-induced drinking/neurophysiology. Representative traces of head location within the conditioning box on drinking trials **(B)** and non-drinking trials **(C)** from a single session in a Wistar rat given alcohol solution. Illustrations at the top of all figure panels in **(D1-4)** illustrate the timecourse of stimuli presentation on each trial. Two seconds of ‘baseline’ data precede the start of each trial, in which a light was illuminated for 2 seconds on one side of the two-sided chamber. A one second ‘delay’ in which no stimuli were activated bridged the light cue and the initiation of sipper movement into the chamber. Sipper movement is represented by the two gray arrows, with the first arrow indicating sipper entry, and the second arrow indicating sipper removal. Fluid was readily available (only on the chamber side cued by the light) between the end of the sipper motor entry (first arrow) and the start of the sipper motor removal (second arrow). **(D1-4)** Mean (±SEM) log-transformed head movement speed changed significantly over time on drinking trials compared to non-drinking trials in P and Wistar rats on both alcohol and water sessions. Green bars denote drinking vs non-drinking trial differences (FDR-corrected rank sum tests; p’s<.05).

### Stereotaxic Surgery and Behavioral Electrophysiology

Following the 15 day acquisition/maintenance of 2CAP, a group of Wistar and P rats with matched alcohol consumption history were selected for electrophysiological experiments, and were unilaterally implanted with multi-tetrode arrays in the mPFC (Linsenbardt and Lapish, 2015). This matching was a critical design feature of these experiments for 3 reasons. First, the use of rats that will reliably consume/self-administer excessive amounts of alcohol under limited access conditions is a prerequisite to identifying how such alcohol consumption alters neurophysiological processes - it is not possible to assess the effects of alcohol consumption in populations that do not drink alcohol. Second, matching for alcohol consumption ensured that any observed differences in physiology were not simply due to differences in alcohol experience. Finally, these matched populations of rodents are directly comparable to human studies in which groups of family history positive and family history negative individuals are matched for drinking history (Kareken et al., 2010).

After a full recovery from surgery, animals were given a period of one week of habituation/acclimation prior to electrophysiological recordings. Animals were habituated to the handling required for incremental lowering of tetrodes prior to the task, and also to navigating the 2CAP environment with the tether connecting the implanted electrode array to the recording hardware. After this habituation period, ≈3 days of 2CAP reinforced with 10% alcohol (v/v) were conducted while electrophysiology was recorded using a 96 channel electrophysiological recording system (Neuralynx, Bozeman, MT). Animals were then given ≈3 water sessions where the sippers contained water. Electrodes were lowered 50-100µm prior to each recording session to collect data from new neuronal ensembles. Following the completion of behavioral testing and electrophysiological recordings, placements were verified via histology (reported in Linsenbardt and Lapish, 2015).

### Video Tracking

One video camera was used in conjunction with ANY-maze software (Wood Dale, IL) to track the head location of animals while they performed the task, and another was used to record high-definition video and audio to identify trials where animals ultimately consumed fluid (drinking trials; Figure 1B) or did not (non-drinking trials; Figure 1C). Digital XY coordinates were converted to voltage and fed directly into electrophysiology hardware where they were recorded in parallel to neuronal activity. Raw tracking values were used to plot the location of the animals within the conditioning apparatus.

### Experimental Design and Statistical Analysis

#### Behavioral statistics

Detailed behavioral results for animals used in electrophysiology studies were recently described (Linsenbardt and Lapish, 2015). Behavioral analyses for the current work were primarily focused on time-locked changes in locomotor behavior in response to the various task stimuli. We were particularly interested in determining if there were differences in behavior between trials in which animals ultimately drank fluid, or did not, as these differences may be related to (or mediated by) computations in the PFC encoding drinking decisions. Head movement speed was positively skewed, so it was first log transformed to normalize. We next evaluated differences between movement speed on a bin-by-bin basis using rank-sum tests, which were followed by Benjamini Hochberg FDR correction for multiple comparisons.

#### General electrophysiology statistics

The results of firing rate over the course of the entire 2CAP sessions for electrophysiology studies were recently described (Linsenbardt and Lapish, 2015). The primary goal herein was to evaluate *cue-induced* alterations in neural activity, which was not evaluated previously. Peri-stimulus time histograms (PSTHs) were created by aligning binned (100ms) spike trains for each neuron to the onset of each trial. PSTHs were smoothed using a Gaussian function with a standard deviation of 300 milliseconds, and softmax normalized to avoid being biased by high firing rate neurons by dividing the firing rate of each neuron by its maximum variance (Ames et al., 2014).

#### Stimuli/Task Responsiveness of Individual Neurons

A signal-to-noise statistic (d-prime, d′) was used to quantify the degree to which each neurons activity changed in response to the task stimuli compared to pre-task (baseline) activity as well as chance (surrogate testing); binned (100ms) spike trains were not transformed or normalized in any way prior to these analyses. Individual neurons were evaluated for the degree of responsiveness using d′ (Gale and Perkel, 2010, Barr et al., 2010). Specifically, d′ was calculated by dividing the absolute values of the mean difference between firing rate during the baseline epoch and the rest of trial by the square root of the sum of their squared deviations. To evaluate the significance of the d′ values, surrogate data were created by taking each neurons spike train and randomly shuffling it 500 times. d′ was then determined for each of the 500 randomly shuffled spike trains and these values were used to compute a 95% confidence interval of the null distribution for each neuron.

#### Mutual Information of individual neurons

Following d′ analyses, we next used mutual information (an information theoretic statistical approach (Cover and Thomas, 2005) to precisely quantify the total *amount* of information encoded by each neuron. This approach is preferable to other parametric statistical analysis of firing rate, as firing rate distributions are highly non-normal (Timme and Lapish, 2018). We focused these analyses on two categorical domains – the amount of information encoding real trials vs null trials (collectively referred to as trial-encoding), and the amount of information encoding drinking trials vs non-drinking trials (collectively referred to as drink-encoding). Null trials were constructed from periods of the neural recording that were randomly selected from the inter-trial interval such that full null trials did not overlap real trials at any time. The detailed statistical and mathematical procedures involved in these analyses are provided as a supplement.

#### Principle component analysis (PCA)

PCA was conducted to evaluate the predominant population-level firing rate dynamics. PCA is commonly used as a dimensionality reduction tool that requires minimal assumptions of the data (Cunningham and Yu, 2014). A single ‘omnibus’ PCA was conducted on a matrix containing all data for all groups so that every possible comparison could be made statistically. This matrix included ensemble activity on drinking trials, non-drinking trials, and equally sized, randomly sampled data vectors (previously described null trials).

#### Neural population State-Space (SS) analyses

For state-space analyses, neural population activity was projected onto the first 3 PCs of PCA space. These analyses allowed us to determine the time course of alterations in the pattern of firing rate. Similar patterns of population activity reside close to each other in 3-dimensional space, and when different are further apart. Differences in distance between 3-dimensional population activity vectors were evaluated on a bin-by-bin basis via Euclidean distance analyses (Ames et al., 2014). The mean distance between each trial and every other trial in that comparison type were made (for example drinking trial 1 vs all null trials, drinking trial 2 vs all null trials, etc.), and the mean and variance of the (non-redundant) distances were used for plotting and statistical analyses. We were specifically interested in differences between drinking and non-drinking trials (vs null trials), and therefore evaluated Euclidean distance between these groups and null trials on a bin by bin bases using Benjamini Hochberg FDR-corrected rank-sum testing.

## Results

### Movement dissociates drinking versus non-drinking trials during fluid availability but not during stimulus (DS) presentation

To assess the neural dynamics of alcohol-associated cues within mPFC, extracellular electrophysiological activity was obtained from ensembles of neurons during performance of an alcohol-drinking task in Wistar and P rats matched for alcohol history (Linsenbardt and Lapish, 2015, McCane et al., 2014). Neural recordings were performed in well-trained animals that had > 7 weeks of prior alcohol experience. Subsequent recordings were made using identical procedures, except the alcohol solution was replaced with water. The layout of the conditioning apparatus (Figure 1A), as well as representative video tracking data on drinking (Figure 1B) and non-drinking (Figure 1C) trials are presented in Figure 1. Head movement speed differentiated drinking from non-drinking trials in both rat populations on both alcohol and water sessions, primarily (or exclusively) during the fluid access epoch (Figure 1D; FDR-corrected rank sum tests; p’s<.05). Differences during fluid access were expected, as drinking required that animals remain in close proximity to the sipper on drinking trials. No differences in movement speed were observed during the DS of drinking versus non-drinking trials, while transient differences were observed from the 2 to 4.5 second period following DS offset (Figure 1D). Furthermore, no differences in the mean number of drinking (one-way ANOVA; F(3,22)=1.02,p=0.41) or non-drinking (one-way ANOVA; F(3,22)=1.05,p=0.40) trials were observed (data not shown). Collectively these data indicate that the behavioral response to the DS was not predictive of a drinking trial.

### Task stimuli elicited differential responsiveness in neurons on drinking trials versus non-drinking trials

To determine if firing rates of individual neurons differed on drinking vs. non-drinking trials, the changes in firing evoked by the presentation of trial-associated stimuli (e.g., DS, sipper) was compared to a baseline period 2 seconds immediately before the trial (Figure 2A). Heterogeneity in the firing rates evoked by task stimuli varied greatly between neurons, with some showing both increases and decreases in firing rate (Figure 2B1), and others displaying only decreases (Figure 2B2) or increases (Figure 2B3). The signal-to-noise statistic d′ was used to identify stimulus responsive neurons, and out of 520 neurons across both groups of rats, 179 (≈34%) displayed statistically significant changes in d′ (Figure 2A+C). Neurons were observed that responded to task-associated stimuli similarly on drinking and non-drinking trials (Figure 2D, purple group). Additionally, subgroups of neurons were then identified that were influenced by task stimuli *only* on drinking trials or non-drinking trials (Figure 2D red and blue groups, respectively). Comparisons of drink-encoding neurons confirmed that non-drinking trial responsive neurons displayed lower responsiveness on drinking trials. The converse was also true; the subgroup of drinking trial responsive neurons displayed lower responsiveness on non-drinking trials (Figure 2D inset). Interestingly, a greater number of responsive neurons were found when drinking status was taken into account compared to when it was ignored (225 vs 179; Figure 2F), with no significant differences in the proportions of neuron response between P and Wistar rats on either alcohol (χ^2^=3.24; p=0.20) or water sessions (χ^2^=2.34; p=0.31; Figure 2E). Thus, mPFC neurons were found that possessed the capacity to encode decisions and/or behaviors associated with drinking/non-drinking trials.

**Figure 2.**
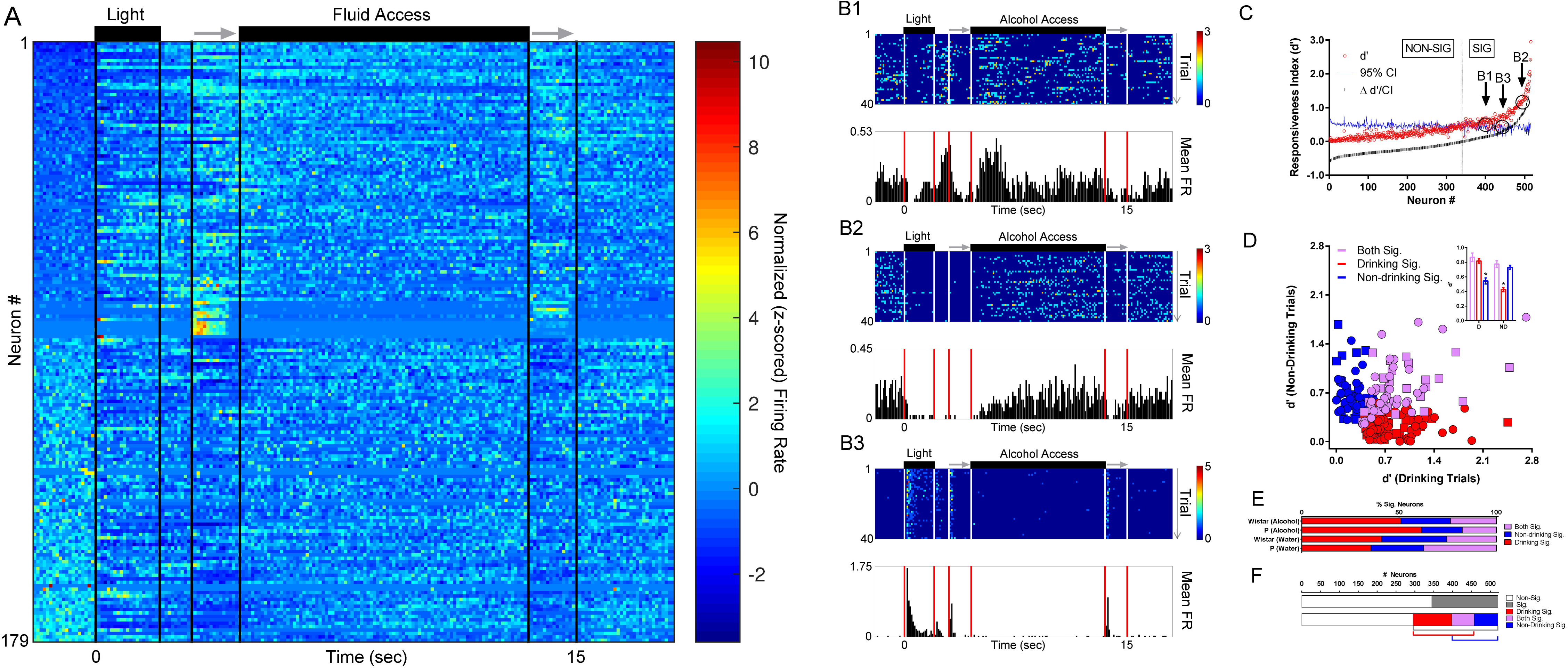
Task stimuli elicited varied responses in neurons on drinking trials versus non-drinking trials, illustrating the capacity to encode/predict future drinking. (**A**) The z-scored timecourse of alterations in firing rate in each of the 179 neurons with significant firing rate alterations (ignoring drinking vs non-drinking status) sorted from lowest baseline firing rate (top) to highest baseline firing rate (bottom). (**B1-B3**) Peri-stimulus time histograms (PSTHs) of 3 representative neurons recorded from a Wistar rat during the same alcohol access session; all displayed significant alterations in firing rate (see panel C). (**C**) Approximately 1/3 of all neurons displayed significant alterations in firing rate vs. baseline as measured by d′ (ignoring drinking vs non-drinking status). (**D**) Significant individual neuron d′ scores on only drinking trials (red), only non-drinking trials (blue), and both drinking and non-drinking trials (purple). Square symbols represent data from Wistar rats and Circle symbols represent data from P rats. The mean of d′ scores on drinking vs non-drinking trials from these subgroups was as expected (inset); drinking-responsive neurons had lower d′ values on non-drinking trials, and non-drinking-responsive neurons had lower d′ values on drinking trials. (Rank-sum tests; KW’s≥50.74; p’s<0.0001). *’s in inset indicate significantly lower d′ scores from other two comparison groups (Dunn’s multiple comparisons post-hoc testing adjusted p’s<.0001). (**E**) The proportion of neurons displaying significant d′ values (drinking/non-drinking/both) were similar between P and Wistar rats (Kruskal-Wallis test; KW=1.54; p=0.71). (**F**) When data were evaluated independently of drinking status (top) a smaller proportion of neurons demonstrated selectivity to presentation of environmental stimuli (≈33%) than when selectivity was assessed taking drinking/non-drinking trials into account (bottom, ≈43%).

### P rats exhibit diminished drink-encoding

To quantify and compare the *amount* of information encoded by trial-and drink-encoding neurons over time, information theoretic statistical approaches were used. The goal of these analyses were to capture the amount of information encoded in each neuron about the trial-associated stimuli (trial-encoding) and if the neural firing rates dissociated drinking/non-drinking trials (drink-encoding). Additionally, these analyses focused on drink-encoding that occurred *prior* to fluid availability (the 0 - 4.5 second interval), as this time interval was expected to contain cue-elicited encoding of the intention to drink or abstain. In addition to quantifying the amount of information using mutual information (MI), these analyses captured different encoding strategies (e.g., firing rate increases or decreases) at each time bin during a trial (e.g., encoding during the DS vs. encoding during access).

Examples of trial-encoding neurons are plotted in Figure 3A1-3. There was marked heterogeneity in trial-encoding. The neurons in Figure 3A1+A3 encoded trial stimuli with increases in firing rate, whereas the neuron in Figure 3A2 did so with decreases in firing rate. The neurons in Figure 3A1+A3 displayed differences from one another in the encoding of the sipper retracting. Additionally, the neurons in Figure 3A2+A3 encoded both visual (light) and auditory stimuli (sipper entry), compared to the neurons in Figure 3A1 which primarily encoded visual (DS) stimuli. Collectively, the neurons recorded from Wistar’s exhibited stronger trial-encoding than P’s during alcohol sessions (FDR-corrected rank sum tests; p’s<.05; Figure 3B), whereas no differences were observed in trial-encoding during water sessions (FDR-corrected rank sum tests; p’s<.05; Figure 3C).

**Figure 3.**
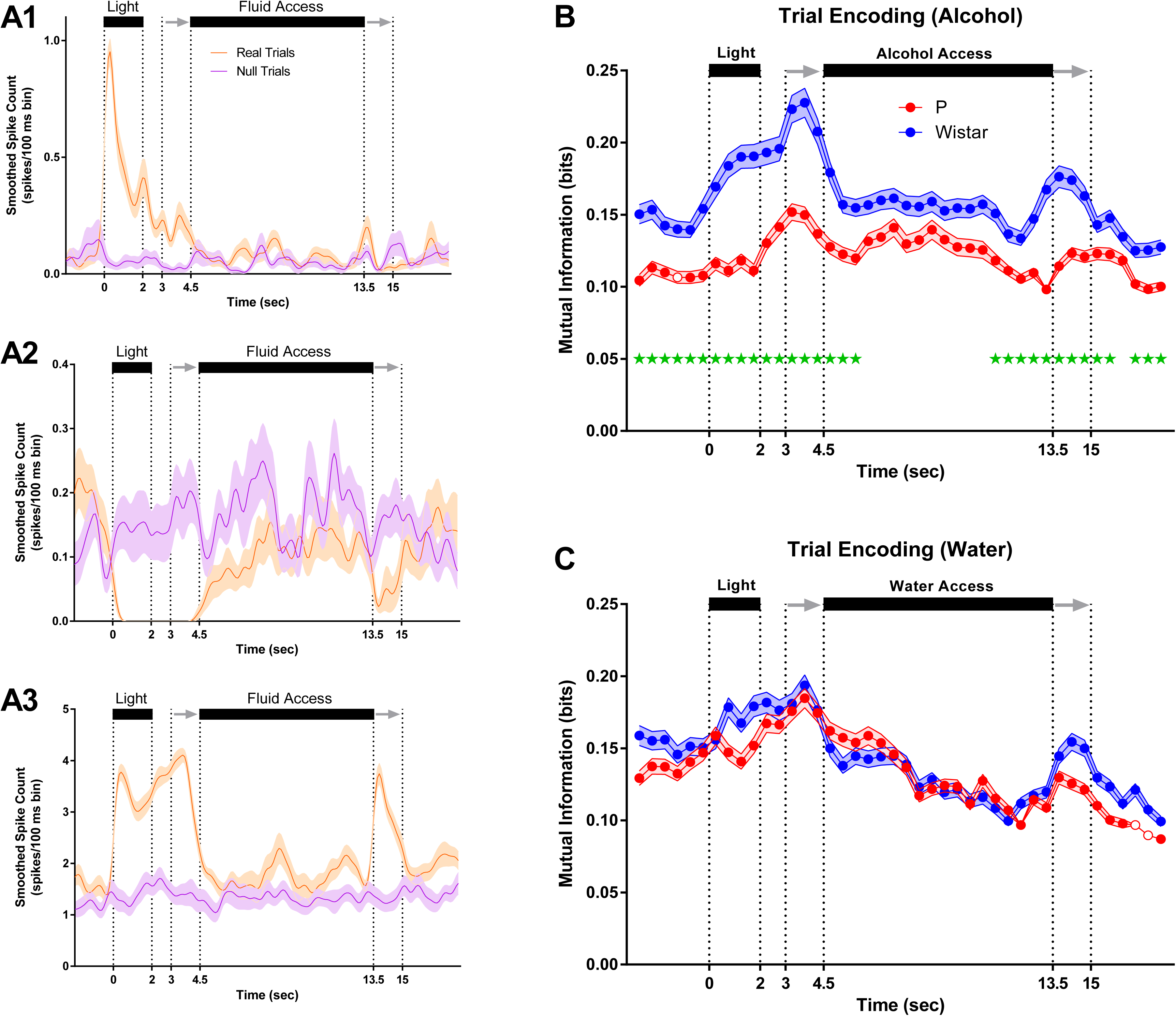
P rats exhibit blunted trial encoding during alcohol sessions. (**A1-A3**) Mean firing rate of 3 representative trial encoding neurons. Neurons in **A1+A3** (Wistar/Alcohol and P/Water) encoded trial stimuli with increases in firing rate, whereas neuron in **A2** (Wistar/Alcohol) did so with decreases in firing rate. Neurons displayed significant heterogeneity in the magnitude and location of trial encoding. For example, neurons in **A1+A3** displayed differences in the encoding of the sipper retracting. Also, **A2+A3** encode both visual and auditory stimuli. On average, Wistar neurons encoded more information about trial stimuli than P during alcohol sessions **(B)**, whereas no differences were observed between P and Wistar during water sessions **(C)**. Data represent weighted mean ± standard error of the weighted mean. Green *’s represent FDR corrected differences between P and Wistar (p < 0.01). Open circles represent time bins where the ensemble of neurons did not produce significant encoding.

Examples of drink-encoding neurons are plotted in Figure 4A1-3. As with trial-encoding, neurons displayed heterogeneity in the magnitude and location of drink-encoding. The neurons in Figure 4A1+A3 encoded drinking intent (pre-fluid availability drink-encoding), whereas the neuron in Figure 4A2 encoded drinking only following fluid availability. The neurons in Figure 4A1-A3 displayed differences from one another in the encoding of drinking during/following fluid removal. Collectively, neurons recorded from Wistar rats encoded more information than P’s about drinking/non-drinking trials prior to alcohol access vs P rats (FDR-corrected rank sum tests; p’s<.05; Figure 4B), which may indicate that the mPFC of Wistar rats performed computations associated with subsequent drinking; such as the intention to drink. In contrast, there were little to no differences in drink-encoding across rat populations prior to water availability (FDR-corrected rank sum tests; p’s<.05; Figure 4C).

**Figure 4.**
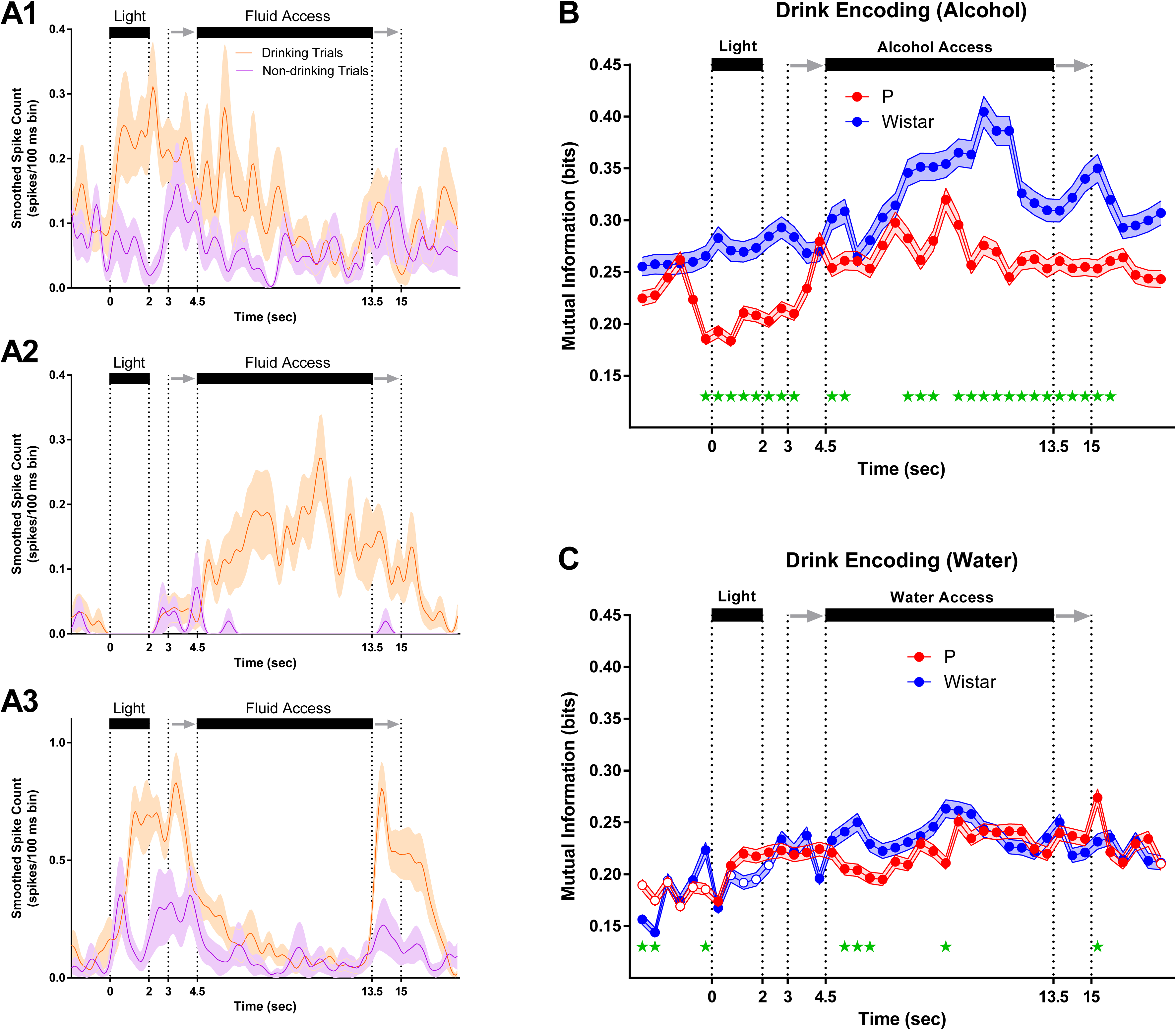
P rats exhibit diminished drink encoding during alcohol sessions. (**A1-A3**) Mean firing rate of 3 representative drink encoding neurons. Neurons in **A1+A3** (Wistar/Water and Wistar/Alcohol) encoded drinking intent (pre-fluid availability drink encoding), whereas neuron in **A2** (P/Alcohol) encodes drinking only during fluid availability. As with trial encoding, neurons displayed significant heterogeneity in the magnitude and location of drink encoding. For example, neurons in **A1-A3** displayed differences in the encoding of drinking during/following fluid removal. On average, Wistar neurons encoded more information about drinking/non-drinking than P during alcohol sessions **(B)**, whereas inconsistent/transient differences were observed between P and Wistar during water sessions **(C)**. Data represent weighted mean ± standard error of the weighted mean. Green *’s represent FDR corrected differences between P and Wistar (p < 0.01). Open circles represent time bins where the ensemble of neurons did not produce significant encoding.

### Neural activity patterns in populations of mPFC neurons reflect the intention to drink in Wistar, but not P, rats

In order to determine if differences in information encoding observed at the single neuron level were maintained at the population level, state-space analyses were performed to quantify how neural activity patterns, captured via principle components, evolved throughout a trial. To quantify the evolution of neural trajectories, Euclidean distance to a corresponding time bin of a null trial was computed for drinking, non-drinking, and null trials (note: a given null trial was compared to all other null trials to compute distance). Euclidean Distance was calculated from a multidimensional space that was defined by the first 3 principle components. Larger values of Euclidean distance correspond to larger differences in neural activity patterns, which indicate that the predominant patterns of neural firing were different for two comparisons (Figure 5A). Supplemental Videos1-4 for each comparison group are provided to illustrate the evolution of neural trajectories over time for each trial. During alcohol sessions, alcohol-associated cues elicited neural activity patterns that diverged prior to the availability of alcohol when drinking-versus non-drinking trials were compared in Wistar (FDR-corrected rank sum tests; p’s<.05; Figure 5B), but not P rats (FDR-corrected rank sum tests; p’s<.05; Figure 5C). In other words, the temporal evolution of neural activity patterns in Wistar rats in response to alcohol-associated cues were predictive of future drinking/non-drinking trials, whereas the neural activity patterns in P rats were not. Additionally, during water sessions, population activity only briefly differentiated drinking trials from non-drinking trials in Wistar (FDR-corrected rank sum tests; p’s<.05; Figure 5D), and failed entirely in P rats (FDR-corrected rank sum tests; p’s<.05; Figure 5E). In contrast, there were large differences between P and Wistar in cue/task-elicited population activity. Specifically, on water drinking trials, P rats displayed greater alterations in neural activity patterns vs Wistar rats (FDR-corrected rank sum tests; p’s<.05; Figure 6A). Thus, in P rats, the mPFC was biased toward encoding alcohol drinking during alcohol consumption, whereas in Wistar rats, encoding of the intention to drink alcohol *and* alcohol drinking was present. Therefore, converging evidence suggests that the encoding of alcohol drinking intent is impaired in the mPFC of P rats, which may contribute to the predisposition for excessive alcohol consumption.

**Figure 5.**
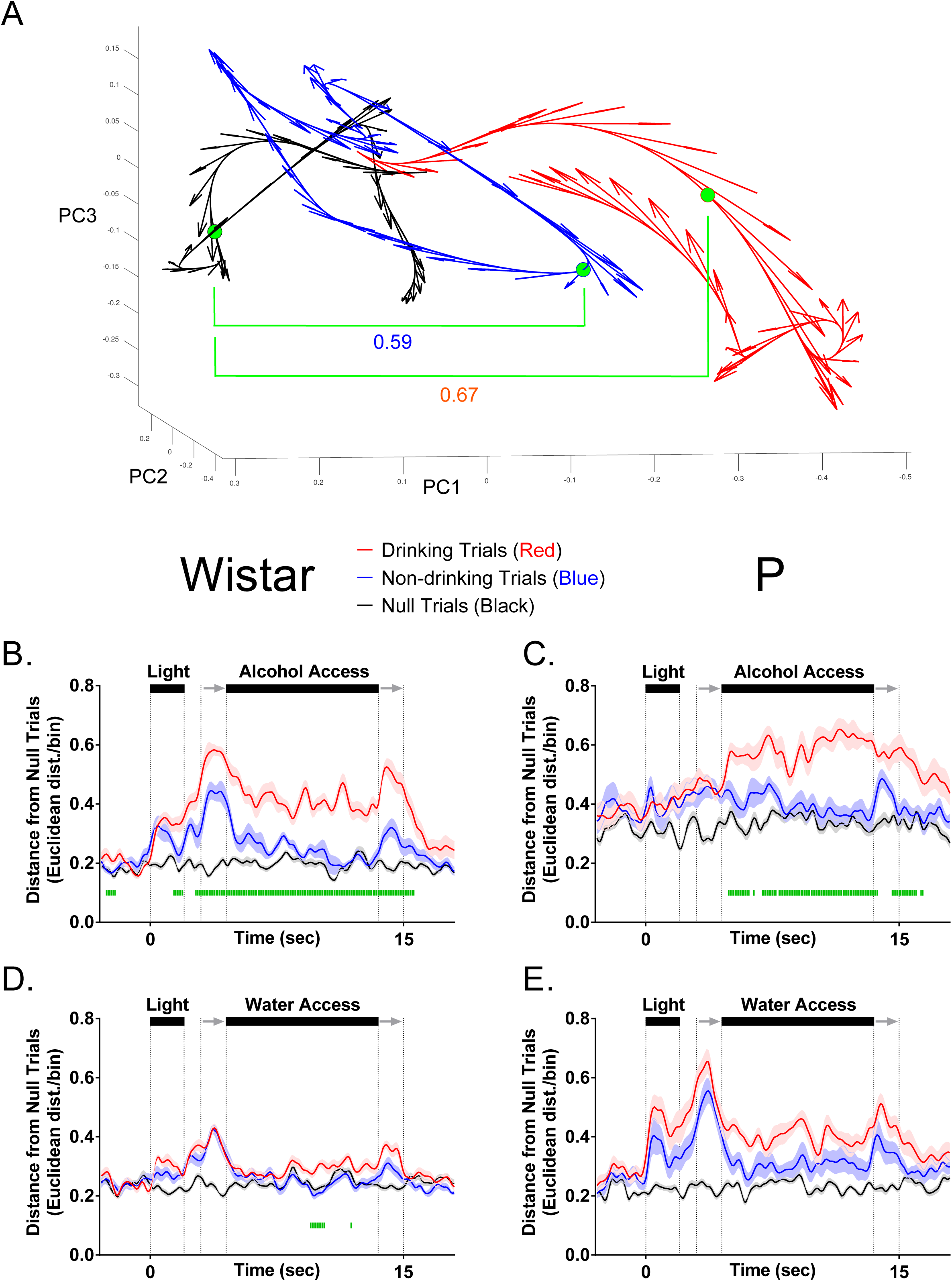
mPFC neural activity patterns reflect the intention to drink alcohol in Wistar, but not P, rats. **(A)** Illustrates neural trajectories in 3-dimensional Euclidean space on a single drinking (red), non-drinking (blue), and null trial (black). Filled green circles indicate the same time bin across each of the conditions, with the Euclidean distance between drinking (0.67) and non-drinking (0.59) trials from null used for statistical analyses in **B-E**. **(B)** Populations of neurons in Wistar rats on alcohol access sessions encoded the intent to drink or not drink – differences in the pattern of firing between drinking/non-drinking trials were observed prior to alcohol access. **(C)** Populations of neurons in P rats on alcohol access sessions encoded drinking/non-drinking, but did not encode alcohol drinking intent. **(D)** Populations of neurons in Wistar only transiently encoded water drinking. **(E)** Populations of neurons in P failed to encode water drinking or water drinking intent. Data are presented as mean ±SEM. Green |’s represent FDR corrected differences in Euclidean Distance between drinking and non-drinking trials (p < 0.05).

**Figure 6.**
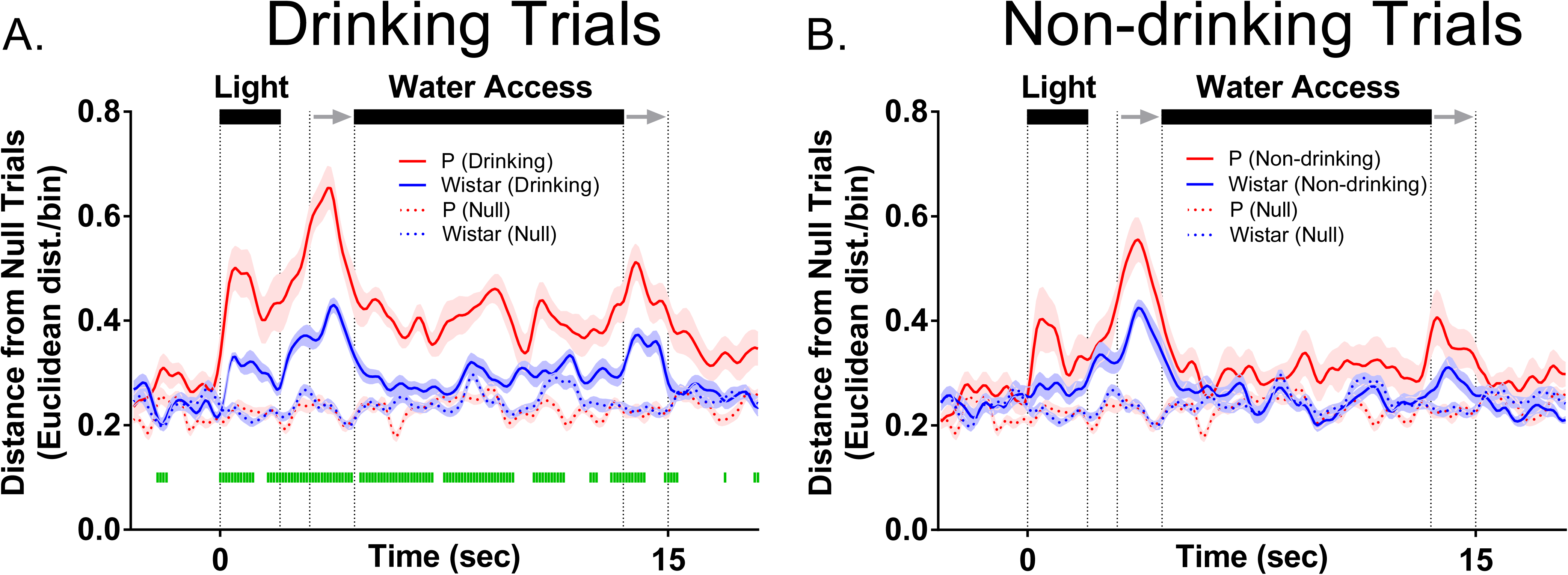
mPFC neural activity patterns more robustly encode alcohol-associated stimuli than Wistar during water sessions. Data presented in this figure are identical to those found in figure 5D+E, and are presented here to illustrate P vs. Wistar differences. **(A)** On drinking trials during water sessions, population of neurons in P rats better encoded alcohol-associated task/stimuli than Wistar rats, whereas there were no differences in encoding of task/stimuli between P and Wistar on non-drinking (water) trials **(B).** Data are presented as mean ±SEM. Green |’s represent FDR corrected differences between P and Wistar (p < 0.05).

## Discussion

The goal of the current study was to determine if the intent to drink alcohol was encoded by populations of neurons in the rodent mPFC, and if such encoding was influenced by a family history of alcohol drinking. Task-stimuli-evoked changes in neural activity were observed in mPFC of both strains of rats for either reinforcer (Figure 2). Contrary to our hypothesis, during alcohol sessions, patterns of neural activity at both the single neuron and population levels more robustly disambiguated drinking from non-drinking trials in Wistar rats. Importantly, these differences were observed *prior to the availability of alcohol,* possibly reflecting the intent to drink (Figures 4B+ 5B). Additionally, during alcohol sessions, enhanced trial-encoding was observed in Wistar rats (Figure 3B), whereas during water sessions, task-stimuli-evoked changes in neural population activity was larger in P rats (Figure6A). Collectively, these data suggest that differences in family history of excessive drinking may alter the computations performed by mPFC that control alcohol drinking, either directly, or as a consequence of an interaction between inherited/genetic differences and moderate (but similar) alcohol history.

### In water sessions, P rats more robustly encode alcohol-associated stimuli

P rats are an extremely well-validated transgenerational rodent model of AUD (McBride and Li, 1998, McBride et al., 2014, Bell et al., 2014, Lumeng and Li, 1986, Gatto et al., 1987, Stewart et al., 1991, Kampov-Polevoy et al., 2000, Waller et al., 1982). One feature that sets these animals apart from other rodent populations that willingly consume alcohol, is their robust seeking phenotype (Czachowski and Samson, 2002). Given this, it was surprising to find that during alcohol seeking/consumption, trial-encoding was weaker in P’s (Figure 3B). Weaker trial-encoding in P’s did not result in an opportunity cost, as there were no differences in the number of drinking trials or volume of fluid consumed between Wistar and P rats. However, differences in drinking were intentionally minimized across Wistars and P’s, as each animal was selected to control for differences in behavior and, especially, history of alcohol intake (Linsenbardt and Lapish, 2015). Since differences in trial-encoding were not associated with increased seeking or drinking when reinforced with alcohol, it does not likely reflect the motivational salience of the stimuli or more basic features of the stimuli such as its perceived intensity or information required to locate/time the delivery of the reinforcer.

Our previous work demonstrated that P rats are slower to extinguish 2CAP (Linsenbardt and Lapish, 2015). This suggests that alcohol-associated stimuli retain their motivational properties in P rats when alcohol is not available. Consistent with this hypothesis, the only comparison where P’s exhibited stronger encoding than Wistars to cues preceding fluid availability was during water sessions at the neural population level (Figure 6A). This observation may reflect the conflict-driven recruitment of mPFC in response to the violation of the previously acquired association between trial-associated stimuli and alcohol experience. Alternatively, enhanced stimuli-encoding during water sessions may reflect the ‘cached’ value of alcohol-associated cues based on prior experiences with alcohol/stimuli rather than their current value (Dezfouli and Balleine, 2013, Daw et al., 2005, Doya, 1999, Rangel et al., 2008, Redish et al., 2008). Consistent with this, an enhanced BOLD response to alcohol associated stimuli in those with increased familial risk for an AUD versus those not at risk has only been observed in a similar setting in which alcohol-associated stimuli are presented in the absence of alcohol access/exposure (Kareken et al., 2010).

### Encoding of the intent to drink alcohol in mPFC is diminished in P rats

Modulation of mPFC inputs to the nucleus accumbens core is capable of remediating aversion-resistant drinking (Seif et al., 2015, Seif et al., 2013) thus indicating that a loss of information originating in mPFC is necessary for the expression of devaluation-insensitive drinking. In the current study, the encoding of drinking intent (e.g., drink outcome specific changes in neural activity *prior* to drinking) was diminished in mPFC of P rats compared to Wistar rats. While, these data are the first to provide evidence that mPFC neurons directly encode the intent to consume alcohol, they also indicate this signal is diminished in animals with increased risk of excessive drinking. These data suggest that increased familial risk diminishes the contribution of the mPFC in the decision to seek and drink alcohol. However, there is also substantial evidence for transitions in encoding in subcortical brain regions, such as the striatum. The dorsomedial striatum directly influences alcohol consumption that is still sensitive to devaluation (i.e., not ‘habitual’), whereas the dorsolateral striatum modulates alcohol consumption only after prolonged training in which animals have become insensitive to devaluation (Corbit et al., 2012). Thus, over the course of repeated alcohol drinking experiences, there is a reorganization of the neural circuits that regulate alcohol drinking behavior. Taken together, these studies underscore the need to disambiguate the distinct roles played by alcohol and family history on the neural circuits that regulate devaluation insensitive and/or aversion-resistant drinking; issues not directly addressed in the current studies.

### Summary/Conclusions

Collectively, the data provided herein indicate differences in the role of the mPFC in alcohol consumption between populations with or without increased familial risk of excessive drinking. This finding is characterized by two primary features observed in P rats: 1) The encoding of the decision to drink is blunted during alcohol drinking, and 2) The encoding of stimuli previously associated with alcohol is enhanced during water sessions. The expression of these features was observed in animals that exhibit an inherited risk for excessive drinking, which may reflect underlying differences in neurobiology that facilitate the resistance to extinction of alcohol seeking and/or the transition to aversion-resistant drinking. Identifying strategies to restore the contribution of the mPFC in the intention to drink alcohol and blunt the encoding of alcohol associated cues observed in water sessions may provide effective targets to treat an AUD. Importantly these data further highlight the need to consider inherited/genetic risk factors when developing treatment strategies for AUD’s.

## Supporting information

## Acknowledgments

This work was supported by NIAAA grant #’s AA022268 (DNL), AA025120 (DNL), AA007462 (NMT), AA022821 (CCL), AA023786 (CCL), the ABMRF (CCL), and the Indiana Alcohol Research Center P60AA007611 (D. Kareken). This research was supported in part by Lilly Endowment, Inc., through its support for the Indiana University Pervasive Technology Institute, and in part by the Indiana METACyt Initiative. The Indiana METACyt Initiative at IU is also supported in part by Lilly Endowment, Inc.

## Author contributions

DNL and CCL designed experiments, DNL conducted experiments, and DNL, CCL, and NMT analyzed data and wrote manuscript.

## Figure Legends

**Supplemental Videos:S1-4.** The top panel in all videos is identical to data found in Figure 6. The bottom panel was generated using DataHigh software (Cowley et al., 2013), and represents the timecourse of neural trajectories over the course of drinking trials (red), non-drinking trials (blue), and null trials (black), in three-dimensional (Euclidean) space. **Suppl_Video_S1** represents data from Wistar rats given alcohol access; **Suppl_Video_S2** represents data from P rats given alcohol access; **Suppl_Video_S3** represents data from Wistar rats given access to Water; **Suppl_Video_S4** represents data from P rats given access to Water.

