## Supplementary material for "Encoding of the Intent to Drink Alcohol by the Prefrontal Cortex is blunted in Rats with a Family History of Excessive Drinking"

**Supplement: Mutual Information**

*Mutual Information of individual neurons*: We began the mutual information calculation by aligning the first 10 drinking, the first 10 non-drinking, and 20 null trials relative to the cue onset (drinking and non-drinking trials) or a randomly chosen time point during the inter-trial interval (null trials). Null trials were constructed from periods of the neural recording that were randomly selected from the inter-trial interval such that full null trials did not overlap real trials at any time. At a given time bin *t* relative to the stimulus onset time and for a given neuron *i*, we constructed a joint discrete probability distribution $p_{i,t}(x,y)$, where *x* was the discretized smoothed spiking rate of the neuron (see above, 100 ms bins relative to stimulus onset) and *y* was either the trial type (real vs. null) or the drink outcome (drinking trials vs. non-drinking trials). The smoothed spiking rate of the neuron was discretized such that the values across trials were ranked and binned into three states (low, medium, or high firing) with equal number of counts (or as close to equal as possible in the event of tied values). The probability was then calculated by dividing the number of joint state observations by the total number of trials. For instance, in the case of drink outcome encoding, for a given neuron and time bin, we may have observed 5 joint states in which the neuron had a low firing rate during drinking trials. In this example, $p_{i,t}(x = \text{low}, y = \text{drinking}) = 5/20 = 0.25$. In the case of stimulus encoding, we might have observed 7 joint states in which the neuron had a high firing rate during real trials. In this example, $p_{i,t}(x = \text{high}, y = \text{real}) = 7/40 = 0.175$. Note that drink encoding only utilized real trials, so only 20 total observations where performed, whereas trial encoding utilized both real and null trials, resulting in 40 total observations.

For each neuron, time bin, and encoding type (drink encoding and trial encoding), we calculated the mutual information using Eq. 1:

$$I_{i,t}(x,y) = \sum_{x,y} p_{i,t}(x,y) log_2 \left(\frac{p_{i,t}(x,y)}{p_{i,t}(x)p_{i,t}(y)}\right) \quad \text{(Eq. 1)}$$

We used base 2 for the logarithm in Eq. 1 to produce mutual information results in units of bits. In Eq. 1, the marginal discrete distributions $p_{i,t}(x)$ and $p_{i,t}(y)$ are found by summing over the other variable:

$$p_{i,t}(x) = \sum_y p_{i,t}(x,y),\ p_{i,t}(y) = \sum_x p_{i,t}(x,y) \quad \text{(Eq. 2)}$$

The mutual information quantifies how much information one variable provides about the other. In this case, if a neuron tends to fire much more frequently on drinking trials than non-drinking trials, for instance, a large mutual information value would result. However, if drinking status and neuron firing rate were unrelated, then a small mutual information value would result. By calculating the mutual information at each time bin for each neuron, we were able to evaluate encoding dynamically throughout the task.

Due to the discrete nature of experimental trials and the fact that mutual information results cannot be lower than 0, noise tends to bias mutual information results upwards (Panzeri et al., 2007, Treves and Panzeri, 1995). To assess the likelihood that a given mutual information result is not simply the result of noise, we calculated a p-value for each mutual information result by randomizing the joint observations 100 times and recalculating the mutual information for these null surrogates. The randomization procedure preserved the marginal distributions. The p-value was then calculated as the proportion of null surrogates with a mutual information result greater than or equal to the observed value in the real data. In the case where all null mutual information values were less than the result from the real data, the p-value was set to 0.005 = 0.5*(1/100) due to the resolution associated with using 100 null surrogates.

Next, to ensure that non-significant mutual information values did not inflate the estimates of standard error, the p-values were used to calculate a weight (w) for each mutual information result via w = $-\log_{10}(p)$. These weights were then normalized by dividing each weight by the sum of all the weights and used to calculate the weighted mean (Eq. 3) and standard error of the weighted mean (Eq. 4) across all relevant neurons (animal strain and liquid type) at a given time bin t.

$$\bar{I}_{w,t}(x,y) = \sum_i w_i I_{i,t}(x,y) \qquad \text{(Eq. 3)}$$

$$SEM_{w,t}(x,y) = \sigma_t \sqrt{\sum_i {w_i}^2} \qquad \text{(Eq. 4)}$$

In Eq. 4, $\sigma_t$ is the standard deviation of the mutual information values across all neurons at the given time bin t. Therefore, large mutual information values that were unlikely to be due to chance received large weights and factored heavily into the weighted mean. In the case where the mutual information results had similar weights, the standard error of the weighted mean approached the standard error of the mean. In the case where the mutual information results were dominated by a few highly weighted values, the standard error of the weighted mean approached the standard deviation.

While the weighting procedure above allowed us to highlight the importance of significant information results, in time bins where few significant information results were observed, an upwards bias in the information results would still be observed. To detect cases where the *ensemble* of information values were not significantly different from null, we also used a KS-test to compare the distribution of real mutual information results to the distribution of mutual information results from null surrogate data used to calculate the individual neuron p-values. This allowed us to assess the time bins for which the entire ensemble of neurons was not significantly different from null data, suggesting the ensemble as a whole was not encoding significant amounts of information (e.g., open circles Figures 3 and 4). We applied a threshold of $p < 0.01$ to all such KS-tests to assess significant ensemble encoding.

Finally, to compare information results between animal populations (P vs Wistar), we used a bootstrap approach to compare the weighted mean mutual information between P and Wistar rats at each time bin. We compared the difference between the weighted mean mutual information values in the real data to the difference weighted mean mutual information results from 10000 randomized trials (identity of P and Wistar neurons randomized preserving number of neurons in each group). The p-value was then calculated as the proportion of randomized trials with differences greater than or equal to the difference in the real data, accounting for the sign of the difference. In the case where all randomized trial difference values were less than the result from the real data, the p-value was set to 0.00005 = 0.5*(1/10000) due to the resolution associated with using 10000 randomized trials. These p-values were then corrected for multiple comparisons across time bins within a given figure using False Discover Rate control (Benjamini and Hochberg, 1995).
